# p53 Regulates Nuclear Architecture to Reduce Carcinogen Sensitivity and Mutagenic Potential

**DOI:** 10.1101/2024.09.14.613067

**Authors:** Devin A. King, Dakota E. McCoy, Soumil Ghosh, Adrian Perdyan, Jakub Mieczkowski, Thierry Douki, Jennifer A. Dionne, Rafael E. Herrera, Ashby J. Morrison

## Abstract

The p53 tumor suppressor is an indispensable regulator of DNA damage responses that accelerates carcinogenesis when mutated. In this report, we uncover a new mechanism by which p53 maintains genomic integrity in the absence of canonical DNA damage response activation. Specifically, loss of p53 dramatically alters chromatin structure at the nuclear periphery, allowing increased transmission of an environmental carcinogen, ultraviolet (UV) radiation, into the nucleus. Genome-wide mapping of UV-induced DNA lesions in p53-deficient primary cells reveals elevated lesion abundance in regions corresponding to locations of high mutation burden in malignant melanomas. These findings uncover a novel role of p53 in the suppression of mutations that contribute to cancer and highlight the critical influence of nuclear architecture in regulating sensitivity to carcinogens.

**One-Sentence Summary:** The p53 tumor suppressor reduces carcinogen sensitivity and mutagenic potential by maintaining nuclear architecture.

---

Environmental carcinogens are a leading cause of mutations that contribute to cancer. The most ubiquitous carcinogen in our natural environment is ultraviolet radiation (UV) in sunlight. Among all cancers, skin cancers are the most frequently diagnosed and have the highest mutational burden in Western societies (*1*).

Defects in repair pathways are known to increase the mutagenic potential of carcinogens. However, much less is known about sensitivity to carcinogens, which is the propensity of different genomic regions to acquire carcinogen-induced DNA lesions - the initial step in mutagenesis. Our previous research demonstrates that sensitivity to UV lesions is regulated by chromatin structure and nuclear architecture (*2*–*4*). Specifically, heterochromatin at the nuclear periphery acquires more UV-induced lesions than euchromatin, and this sensitivity can be altered via inhibition of heterochromatin formation.

Notably, many chromatin regulators are also cancer drivers. For example, several tumor suppressors are transcription factors that alter chromatin structure to regulate the expression of target genes. A classic example is the p53 tumor suppressor that orchestrates gene expression programs in DNA repair, cell cycle, and apoptotic pathways through chromatin modification (*5*). Interestingly, current studies continue to reveal the complexity of p53 tumor suppressor function. New transcriptional targets have been identified, such as lncRNAs and transposons (*6*–*8*). In addition, roles for p53 in enhancer and long-range genomic interactions are emerging (*9*–*11*). However, the influence of p53 on carcinogen sensitivity has not previously been explored. In this report, we investigate novel genome stability functions for the p53 tumor suppressor in the context of carcinogen sensitivity.

## Results

### Loss of p53 increases UV sensitivity without a full canonical DNA damage response

To quantitatively assess acute UV sensitivity profiles in cells, we developed a technique called UV Tag-Seq (<u>U</u>ltra<u>v</u>iolet Lesion <u>Tag</u>)-<u>Seq</u>uencing, which utilizes highly specific DNA repair enzymes and low-input, transposition-based library construction to capture DNA with UV-induced cyclobutane pyrimidine dimers (CPDs) (Fig. 1A) (*12*). UV lesion spike-in standards and unique molecular identifiers (UMIs) facilitate normalization and quantification of lesion abundances between samples.

**Fig. 1.**
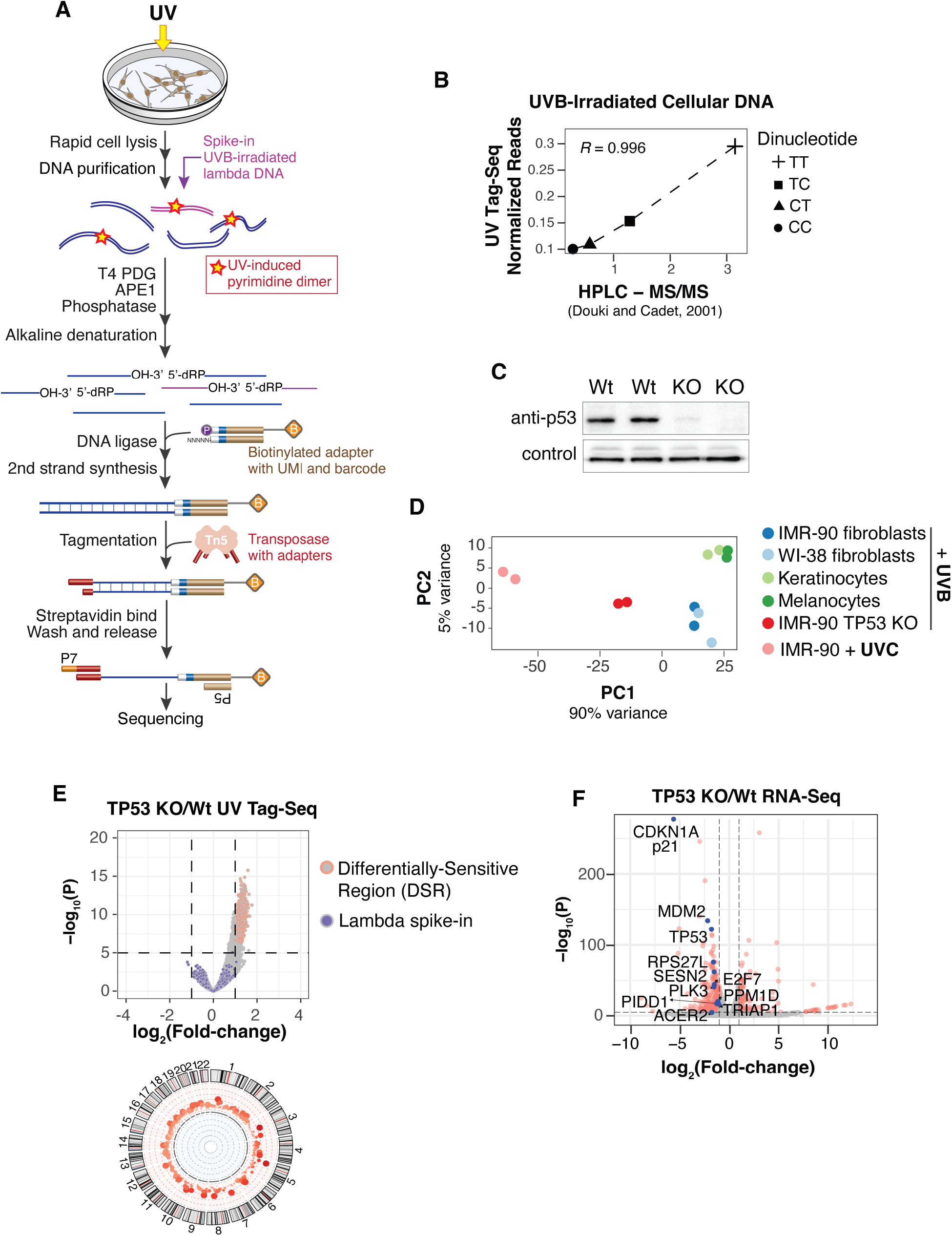
Loss of p53 increases UV sensitivity in the absence of DNA damage response activation. **(A)** Schematic of UV Tag-seq method to quantitatively map UV-induced DNA lesions genome-wide. **(B)** Correlation of UVB-induced dipyrimidines detected by UV Tag-seq compared with bulk quantification by HPLC-MS/MS (*14*). **(C)** Western blot of p53 in wild-type and TP53 knock-out (KO) cells. Technical replicates are shown with loading control (Emerin). **(D)** Principal Component Analysis (PCA) of UV lesions genome-wide in the indicated cells. **(E)** (Top) Volcano plot of differential UV sensitivity for TP53 KO cells compared to IMR-90 wild-type cells treated with 200J/m^2^ UVB. Irradiated lambda DNA used to normalize samples is shown. (Bottom) Circos plot of Differentially Sensitive Regions (DSRs) in autosomes. **(F)** Volcano plot of RNA-seq comparing TP53 KO and IMR-90 wild-type cells. Red dots denote significantly differentially expressed (SDE) genes. Genes in the “DNA damage response, signal transduction by p53 class mediator” gene ontology pathway are highlighted in blue with corresponding names. Non-SDE genes are not highlighted.

We performed UV Tag-Seq on wild-type primary cells after a brief (<10 seconds) exposure to UV followed by immediate cell lysis. The results show that the predominant pattern of lesion accumulation is in heterochromatic regions of the genome (fig. S1A), recapitulating our previous results using an immunoprecipitation-based assay (*2*). The UV Tag-Seq method was further validated by demonstrating that the normalized abundances of pyrimidine dimers (TT>TC>CT/CC) were consistent with a separate study (*10*) that quantified CPDs by mass spectrometry (Fig. 1B and fig. S1B)

UV lesions were then mapped in *TP53* knock-out (TP53 KO) primary cells, created via CRISPR-Cas9 genome editing (Fig. 1C). Also included were primary melanocytes and keratinocytes - cells of origin for melanoma and basal/squamous cell carcinomas, respectively (Fig. 1D). To limit potential cell cycle differences, cells were grown to confluency before irradiation. A physiological dose of 200 J/m2 UVB was used, equivalent to a minimal erythema dose (i.e. skin reddening) for a Caucasian with light brown or olive skin who tans easily (*13*). As before, the duration of UV exposure was brief (<10 seconds), and cells were immediately lysed thereafter.

Intriguingly, the TP53 KO cells exhibited a significant increase in UV sensitivity compared to wild-type cells exposed to the same dose of UVB (Figs. 1D and E). Furthermore, this increase in sensitivity approached that of cells exposed to higher-energy UVC, which is at least 10 times more efficient at producing CPDs than UVB (Fig 1D) (*14, 15*). An overall increase of UV lesions in TP53 KO cells was also confirmed using quantitative mass spectrometry (fig. S1C).

In our experiments, elevated UV sensitivity in TP53 KO cells should not result from deficiencies in a canonical p53-mediated DNA damage response, as the rapid time between UV exposure and cell lysis precludes full activation of damage responses (*16*). However, RNA-Seq analysis demonstrates that a few genes in the damage response pathway, such as CDKN1A/p21, are differentially expressed in TP53 KO cells, even in the absence of damage exposure (Fig. 1F, figs. S2A and B). Thus, while a canonical DNA damage response does not participate in the regulation of UV sensitivity, some components of DNA damage-related responses might be involved.

### Melanoma mutation rates are elevated in p53 UV-sensitive regions

UV-induced DNA lesions are the initial step for the vast majority of mutations in skin cancers, thus the relationship between UV sensitivity and melanoma mutation rate was explored. The aggregated frequency of sunlight-induced C to T mutational signatures in melanomas (*17*) was found to be significantly elevated within TP53 KO DSRs compared to the rest of the genome (Figs. 2A and 2B). In addition, when comparing DSR to non-DSR mutation rates within individual tumors, the majority exhibit increased mutation frequency (Fig. 2C). However, some tumors have either no significant difference or a decrease in DSR vs non-DSR mutation rate. Upon further investigation, this appears to be related to differences in mutagenic signatures and p53 pathway mutation status of these specific tumors (Fig. 2D). Specifically, melanomas originating through non-UV mechanisms, such as acral melanomas (*18*), have reduced or similar mutation rates in TP53 KO DSRs compared to non-DSRs. Extending this analysis to mutation rates in many different tumor types demonstrates that skin cancers have the highest percent difference between TP53 KO DSRs and non-DSRs (Fig. 2E). Conversely, cancers with mutational signatures that are less associated with carcinogens, such as breast and cervical cancer, exhibit relatively low differences in mutation rates between DSRs and non-DSRs.

**Fig. 2.**
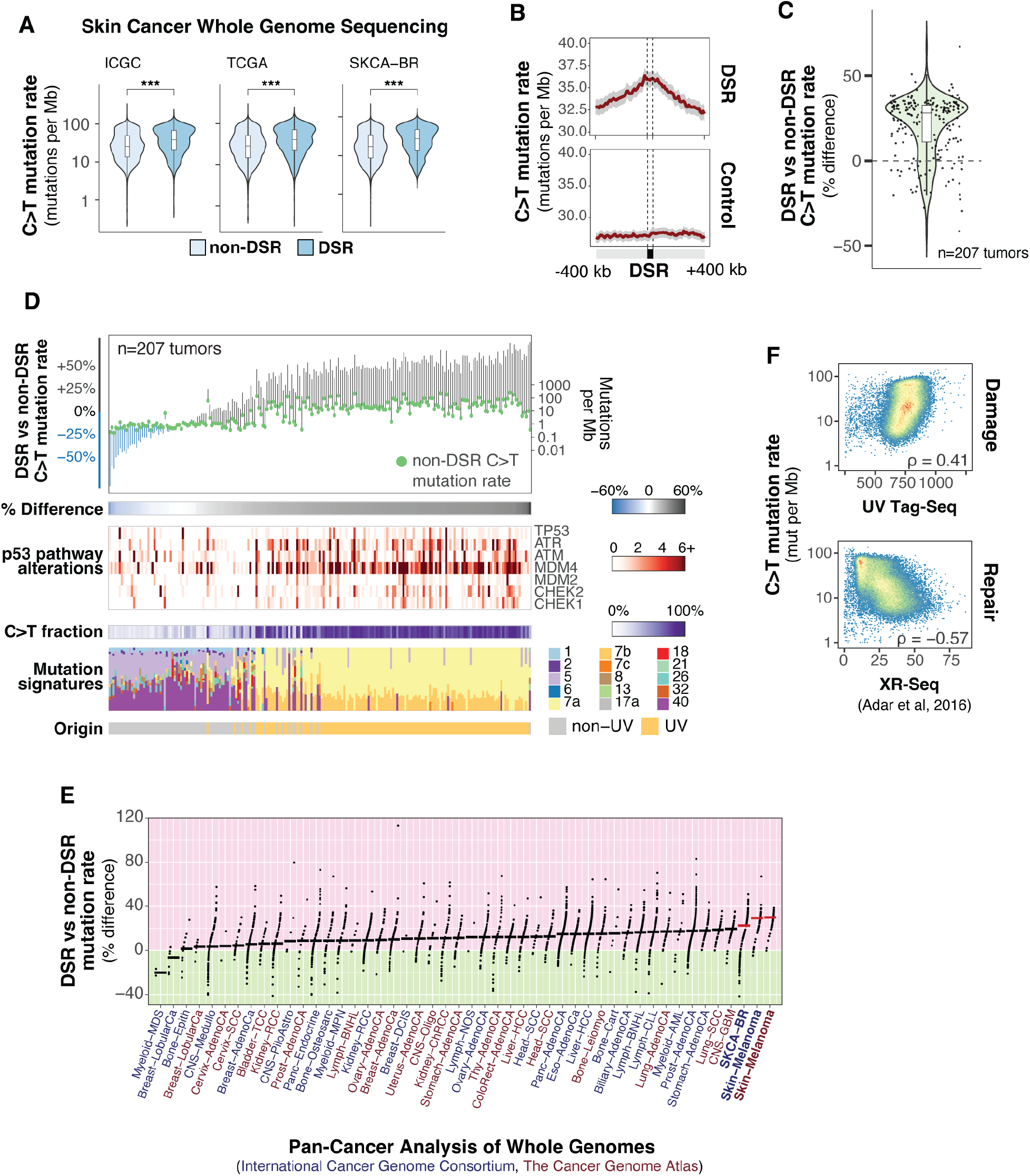
Melanoma mutation rates are elevated in TP53 KO Differentially Sensitive Regions (DSRs). **(A)** C>T mutation rates in TP53 KO DSRs and non-DSRs from whole-genome sequencing studies. ICGC is the International Cancer Genome Consortium, n=70 tumors. TCGA is The Cancer Genome Atlas, n=37 tumors. SKCA-BR is ICGC Skin Adenocarcinoma – Brazil, n=100 tumors. *** indicates Mann-Whitney U test p-value < 0.001. **(B)** Mean mutation rates aggregated in DSRs and randomly sampled regions (control) from samples in (A). Gray bars represent 95% confidence intervals. **(C)** Percent change in DSR vs. non-DSR mutation rate by tumor from samples in (A). **(D)** Tumors described in (A) sorted by percent change in DSR vs. non-DSR mutation rate. COSMIC Mutational Signatures are shown on the bottom. Signature 7 is associated with ultraviolet light exposure. **(E)** 3,022 deep-sequenced tumors from Pan-Cancer Analysis of Whole Genomes (PCAWG) grouped by subproject (anatomical location). Samples are ordered according to median percent change (horizontal bars) in DSR vs. non-DSR mutation rate for each study. ICGC studies are in dark blue and TCGA studies are in dark red. Melanoma studies are bolded with red horizontal bars indicating the median percent change. **(F)** Spearman rank correlation coefficients of UV damage (top panel, UV Tag-Seq) and cumulative nucleotide excision repair (bottom panel, XR-Seq from (*21*)) with C>T mutation rate in tumors shown in (A).

While the formation of UV-induced DNA lesions is the initial step in melanoma mutagenesis, DNA repair is also a critical contributing factor. Indeed, genome-wide analysis of nucleotide excision repair (NER), the primary UV lesion repair pathway (*19*–*21*), in immortalized fibroblasts identified a negative correlation between melanoma mutation rate and repair, while UV sensitivity exhibits a positive correlation (Fig. 2F). Thus, both UV sensitivity and repair deficiency likely contribute to high rates of UV-induced mutation in melanomas.

### p53 regulates UV sensitivity at the nuclear periphery

As chromatin features have previously been found to influence UV sensitivity (*2, 3*), we assessed histone modifications in the top TP53 KO DSRs to investigate the epigenetic context and potential mechanisms of differential sensitivity regulation. These TP53 KO DSRs were found to be enriched in the H3K9me3 heterochromatin mark and depleted in many euchromatic modifications (Fig. 3A). Accordingly, TP53 KO DSRs were enriched in ‘heterochromatin’ and ‘quiescent’ chromatin states (Figs. 3B and C), previously defined by the Roadmap Epigenomics Project (*22*). Heterochromatin is often associated with the nuclear lamina, a filamentous protein structure at the nuclear membrane important for genome organization and membrane structure (*23*). Accordingly, lamin B1 was enriched within TP53 KO DSRs (Fig. 3D). Differential UV sensitivity was then highlighted on a three-dimensional (3D) computational model of the genome in the nucleus (*24*). A striking pattern was observed in that regions with the highest sensitivity in TP53 KO cells were located at the nuclear periphery, while regions closer to the center acquired comparatively fewer UV lesions (Fig. 3E). Collectively, these results demonstrate that the nuclear features associated with increased UV sensitivity in wild-type cells ((*2*), fig. S1A) are further elevated in TP53 KO cells.

**Fig. 3.**
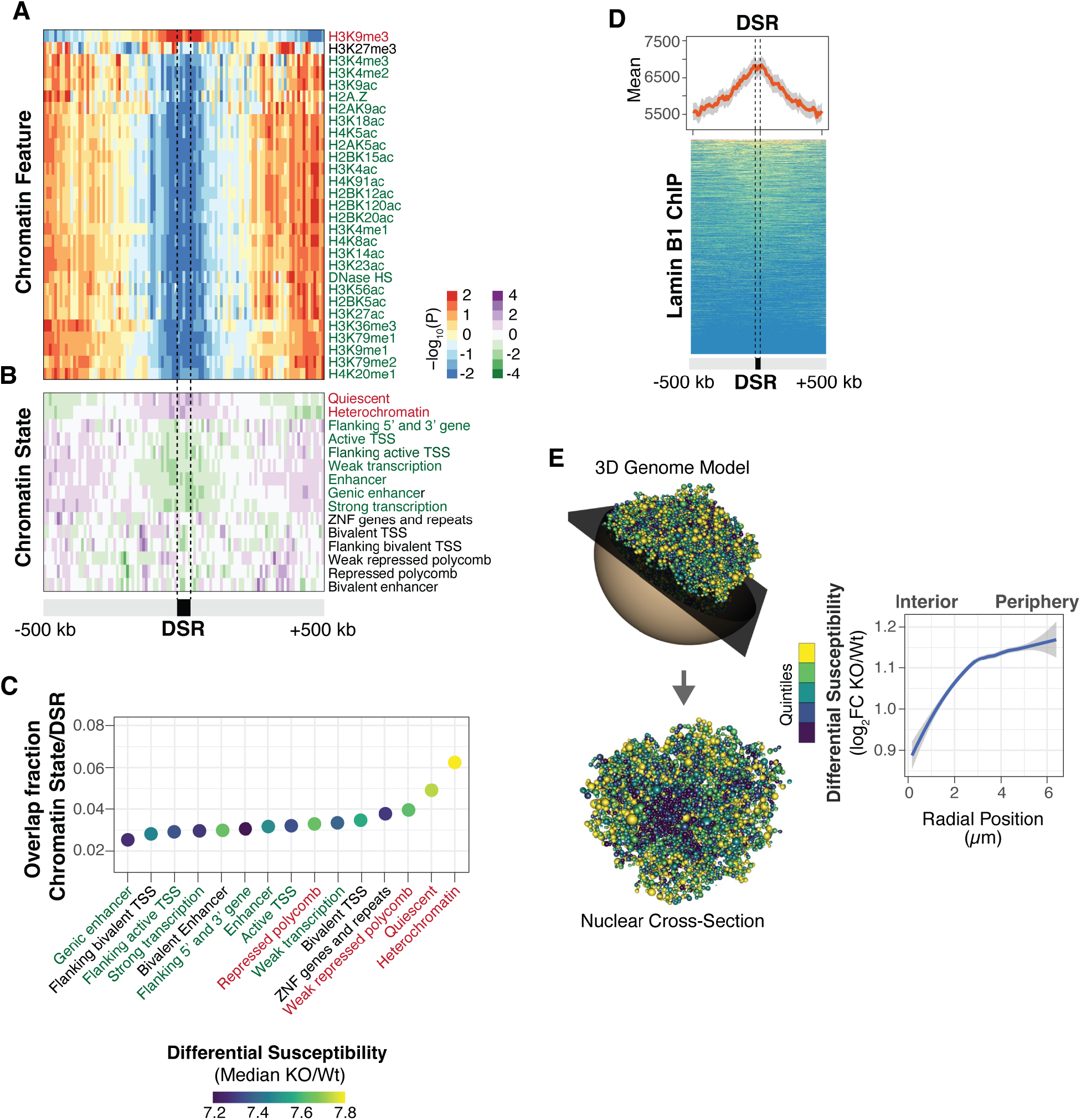
Heterochromatic lamin-associated regions are more UV-sensitive in TP53 KO cells. Enrichment of chromatin features **(A)** and chromatin states from Roadmap Epigenomics Consortium **(B)** in top 5% TP53 KO differential sensitive regions (DSRs). The color of the histone modification or chromatin state name indicates euchromatin (green), heterochromatin (red), or bivalent/mixed (black). **(C)** The dot plot indicates the fraction of overlap between chromatin states and TP53 KO DSRs. The color of the dot represents the magnitude of differential UV sensitivity between TP53 KO and wild-type. **(D)** Composite plot and heatmap of lamin B1 enrichment in TP53 KO DSRs. The line is the mean signal with 95% confidence interval in gray. **(E)** Three-dimensional model of the wild-type genome with TP53KO/wild-type log2 fold-change differential sensitivity colored by quintiles (left) and radial positioning measurements from nuclear center to periphery (right). The model represents the 3D genome of wild-type cells. Each bead in the model represents a Topologically-Associated Domain (TAD) and is assigned specific x, y, and z coordinates.

To examine if TP53 KO DSRs could be directly regulated by p53, the relationship between p53 binding sites and DSRs was assessed in different cell lines from ENCODE (*25*). Analysis of p53 ChIP-seq peaks demonstrates that only a minor portion (<15%) of DSRs overlap TP53 binding sites (fig. S3), suggesting that the majority of TP53 KO DSRs do not appear to be regulated by proximal binding of p53 to target sites.

### TP53 KO cells lack dense chromatin at the nuclear periphery that refracts UV light

Because TP53 KO cells exhibit increased UV sensitivity in genomic regions enriched at the nuclear periphery, we examined various features of the nuclear membrane. Microscopy was first performed with Lamin B1 and DNA staining of confluent wild-type and mutant cells. In the TP53 KO cells, significant differences in nuclear shape, such as excessive membrane curvature, were immediately apparent (Fig. 4A). Automated image analysis of several nuclear morphological features, including shape and symmetry, confirms these abnormalities in TP53 KO nuclei, which are distinct, and more variable compared to wild-type nuclei (Fig. 4B and C). Transmission electron microscopy (TEM) was then performed to examine nuclear structure at higher resolution. The results were striking in that the dense heterochromatin at the nuclear periphery of wild-type nuclei was dramatically reduced in TP53 KO cells (Fig. 4D upper panels). Quantitative image analysis confirms thinner peripheral heterochromatin in TP53 KO nuclei compared to wild-type (Fig. 4E).

**Fig. 4.**
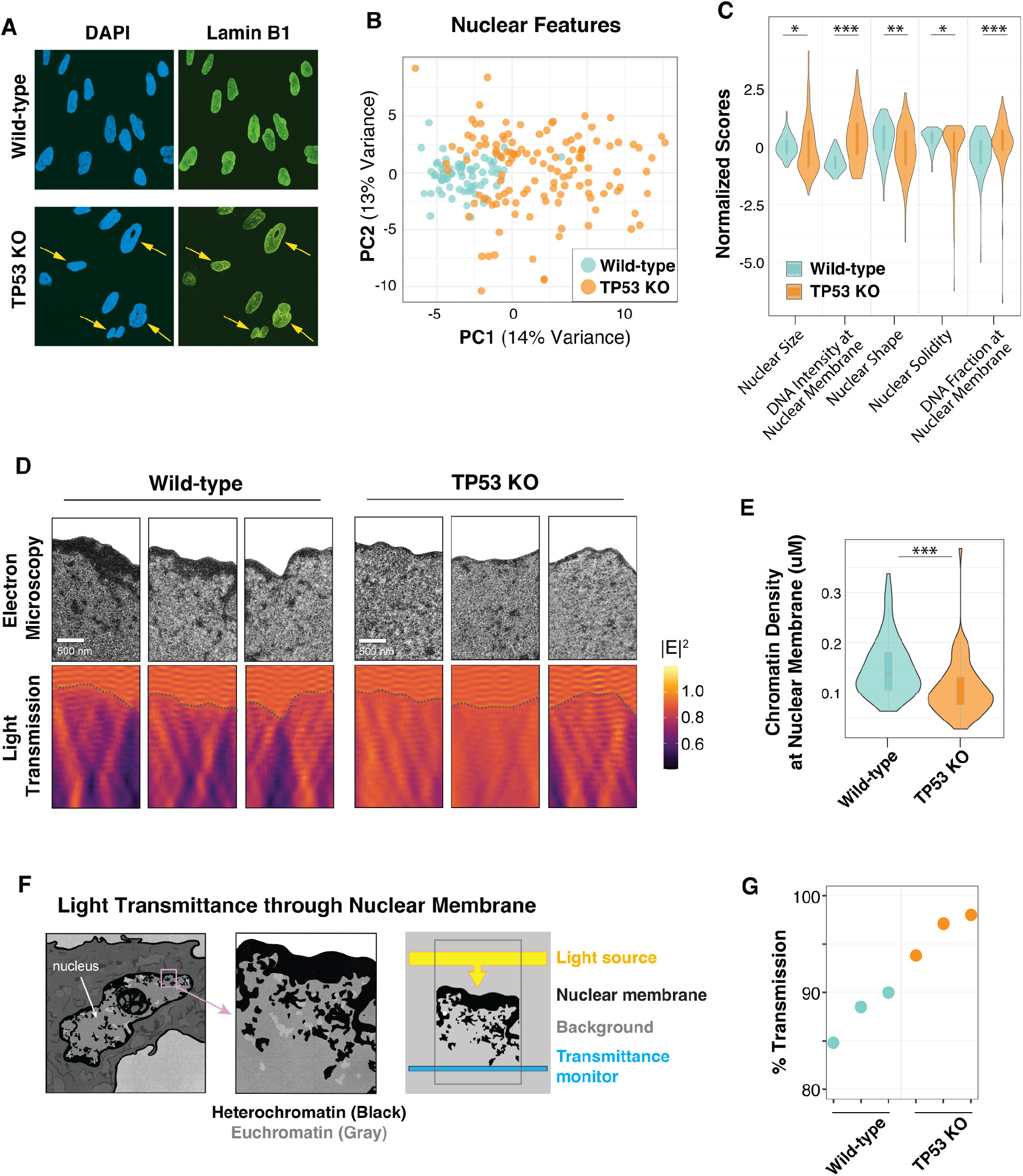
The nuclear membrane of TP53 KO cells exhibits altered morphology and increased light transmission. **(A)** Representative images of lamin B1 and DAPI staining from confluent cells. **(B)** Principal Component Analysis (PCA) of CellProfiler nuclear features that predict wild-type (n=71) and TP53 KO (n=125) cell morphology. **(C)** Representative z-scaled scores for select nuclear features used in the PCA plot in (B). Mann-Whitney U test indicates: * for <0.05; ** for <0.01; *** for <0.001 p-value **(D)** (Top panels) Transmission electron microscopy (TEM) images with the cytoplasm cropped to highlight the nuclear membrane. (Bottom panels) Finite-difference time-domain simulations of light transmission (described in F) for TEM images in top panels. Heatmap indicates the electric field (E^2^). **(E)** Nuclear membrane density measurements of TEM images (n=70) for each cell line. Two sample t-test was indicates: * for <0.05; ** for <0.01; *** for <0.001 p-value. **(F)** Schematic of finite-difference time-domain light transmission simulations using TEM images. Artwork credit to Ana Kimber. **(G)** Measurements of simulated 306 nm UVB light transmission in bottom panels of (D). t-test indicates p < 0.05 significance between cell types.

Chromatin density has previously been shown to alter the optical properties of the nucleus. Specifically, dense heterochromatin is typically more refractive than euchromatin (*26, 27*). We simulated the transmission of light photons through the nuclei of wild-type and TP53 KO cells using two-dimensional full-field electromagnetic simulations (*28*) (Fig. 4F). As proof of principle, a two-dimensional model of different chromatin densities at the nuclear periphery was assessed and found to scale linearly with light transmission (fig. S4). When using actual TEM images from wild-type and TP53 KO cells, the simulated light transmission was higher in the nuclei of cells lacking TP53 (Fig. 4D lower panels, and Fig. 4G).

## Discussion

In this study, we uncover a previously uncharacterized tumor suppressor function for p53 that regulates carcinogen sensitivity and mutagenic potential. Often referred to as the “guardian of the genome”, p53 is the most commonly mutated gene in human tumors and has been extensively studied in DNA damage response pathways (*29*). Results from this study expand p53’s role in genome maintenance pathways and highlight its importance in the regulation of carcinogen sensitivity, independent of DNA damage responses. During carcinogenesis, disruption of p53 function would be expected to increase mutagenic potential by altering both UV sensitivity and repair efficiency.

Loss of p53 dramatically alters chromatin architecture at the nuclear periphery, thereby removing its protective function and increasing the transmission of UV radiation into the nucleus. While heterochromatin at the nuclear periphery is relatively gene-poor, these regions do contain several cancer-driver genes, such as *BRAF* oncogene and *MECOM*, which are both located directly in lamina-associated domains (LADs) (*2*) and are mutated in over 50% and 20% of malignant melanomas, respectively (*30*). Likewise, the *NRAS* oncogene is approximately 11 Kb from a LAD and is mutated in 27% of melanomas. Indeed, many critical drivers of skin cancer are in regions of high sensitivity, thereby increasing their potential contribution to malignant transformation (*2*). Interestingly, a separate study found that depletion of p53 in immortalized primary cells compromises the integrity of the nuclear envelope, resulting in spontaneous rupture (*31*). Furthermore, a high-throughput screen in epithelial cells identified p53-depleted cells as one of the cell types with the highest degree of altered nuclear shape (*32*). p53 levels have also been directly linked to defects in nuclear shape and chromatin density in cancer cells (*33*). These features have clinical significance, as alterations in nuclear morphology can increase membrane flexibility, allowing cells to move through the vasculature and travel to distant metastatic sites. Indeed, nuclear morphology is commonly used by pathologists to determine cancer grade and prognosis (*34*). Results from this study expand our understanding of how nuclear structure defects can contribute to cancer development. Furthermore, these findings raise the possibility that other regulators of nuclear membrane integrity may have roles in carcinogen sensitivity.

The specific mechanisms by which p53 regulates nuclear structure are currently under investigation. The mechanical stability of the nucleus is regulated by several cellular components, such as chromatin, lamins, and the cytoskeleton (*35*). Because of its characterized roles in transcriptional regulation via histone modification, p53 may control nuclear shape by altering the polymer-like stiffness of chromatin (*36*). p53 has also been linked to cytoskeleton dynamics during cancer cell migration and epithelial wound repair (*37, 38*). Thus, p53 has the potential to alter nuclear structure and UV sensitivity through different cellular mechanisms that are not directly linked to genome stability.

Regardless of the mechanism, it is expected that alterations in carcinogen sensitivity are not limited to UV light and may also be relevant to any exogenous genotoxin that can diffuse into the nucleus and react with DNA, including other carcinogens and chemotherapeutic agents. Overall, this study highlights the potential impacts of deregulated chromatin structure and nuclear architecture on genome stability and cancer evolution.

## Acknowledgments

We thank Jon Mulholland at Stanford’s Cell Science Imaging Facility for assistance with the electron microscopy, and José Dinneny’s lab for assistance with the fluorescent imaging. We are grateful for Prof. Andrew Spakowitz’s helpful assistance with the photophysical models. We thank the Stanford Research Computing Center for providing computational resources that contributed to these research results.

## Funding

National Institutes of Health grant R21CA178529 (AJM)

National Institutes of Health grant R21CA171050 (AJM)

National Science Foundation Fellowship GRFP DGE-1656518 (DAK)

Chan Zuckerberg Biohub, San Francisco (JAD)

National Science Foundation PRFB Program grant 2109465 (DEM)

Stanford Science Fellowship (DEM)

NAWA-Polish National Agency for Academic Exchange, Walczak’s Scholarship BPN/WAL/2022/1/00024/U/00001 (AJP)

National Institutes of Health grant S10OD028536 (JWM)

## CRediT author contributions

Conceptualization: AJM, REH

Methodology: REH, DAK, SG, AJM, JAD

Investigation: DAK, REH, AJM, DEM, TD, AP

Software: DAK, SG

Funding acquisition: AJM, DAK, JAD, DEM, AP

Supervision: AJM, REH, JAD, JM

Writing – original draft: AJM

